# A multiscale map of the descending sympathetic innervation of the spleen

**DOI:** 10.64898/2026.11.15.766191

**Authors:** Leonor Remédio, Zachary R. Lewis, Emma D. Thomas, Ines Rodrigues-Vaz, Christine R. Schmidt, Lin T. Tun, Tiffany X Chen, Katherine A. Fancher, Joshua S. Greene, Darren Bertagnolli, Jeff Goldy, Christine Rimorin, Michael Tieu, Trangthanh Cardenas, Amy Torkelson, Junitta Guzman, Rebecca Ferrer, Anish Bhaswanth Chakka, Ana M. Vicente, Shenqin Yao, Kimberly A. Smith, Bosiljka Tasic, Darcy S. Peterka, Rui M. Costa

**Affiliations:** Allen Institute, Brain Science, Neurobiology of Action and Neural Dynamics, Seattle, WA United States; Zuckerman Mind Brain Behavior Institute, Columbia University, New York, NY United States

## Abstract

Brain regulation of visceral organ function depends on descending neural circuits. The spleen is the largest secondary lymphoid organ, and studies indicate that descending neural activity can alter splenic immune responses. Here, we combined retrograde tracing, single-nucleus RNA sequencing, spinal cord and brain reconstruction, and glycoprotein-complemented rabies tracing to map the sympathetic circuits associated with these organs in mice. Catecholaminergic fibers densely innervated the spleen and pancreas, and retrograde labeling identified largely distinct populations of postganglionic neurons within the celiac-superior mesenteric ganglion complex (CMG). Spleen-projecting neurons were enriched in suprarenal regions, whereas pancreas-projecting neurons were distributed more broadly. Single nucleus transcriptomics resolved three CMG neuron types and a Cre line marking the Rxfp1+ neurons labeled spleen-projecting neurons. CMG-projecting sympathetic preganglionic neurons occupied the intermediolateral cell column, intercalated nucleus, and spinal nucleus of the thoracic spinal cord. A tracing strategy designed to identify neurons upstream of these preganglionic populations labeled neurons in the medulla, pons, midbrain, hypothalamus, and several areas of the cortex including motor cortex. Motor-cortex axons formed vGlut1-positive boutons in close apposition to retrogradely labeled CMG-projecting neurons, providing anatomical support for a descending cortical route to sympathetic spinal circuits. These data provide an integrated anatomical and transcriptomic description of the descending circuits innervating the spleen and suggest that motor cortex is an effector cortex that controls more than movement through its projections to the spinal cord.

## INTRODUCTION

Coordinated responses to infection and tissue injury involve communication among immune organs, circulating cells, cytokines, and the nervous system. The spleen, the largest secondary lymphoid organ, is densely innervated, and previous studies indicate that descending neural activity can alter splenic immune responses^1–5^. However, the organization of the descending circuits linking the brain, spinal cord, sympathetic ganglia, and spleen remain incompletely resolved.

The spleen contributes to immunity, hematopoiesis, and inflammation. Autonomic innervation of the spleen has been described across vertebrate species^6–11^. Within the spleen, sympathetic fibers accompany blood vessels, particularly arterioles, and extend into the parenchyma. In rodents, sympathetic varicosities lie in close apposition to lymphocytes, which express adrenergic receptors^1,2,12^. This conserved anatomical arrangement provides a substrate through which sympathetic activity could influence vascular and immune functions.

Manipulating splenic nerve activity affects both vascular tone and immune function. Stimulation of the splenic nerve, or application of adrenaline or noradrenaline, evokes splenic vasoconstriction in humans, pigs, and rats^13–15^. In rodents, the inflammatory reflex links detection of inflammatory signals to increased vagal and splenic nerve activity and suppression of pro-inflammatory cytokine production. In one proposed mechanism, noradrenaline released from sympathetic terminals promotes acetylcholine release from β2-adrenergic receptor-expressing T cells; acetylcholine then acts on α7 nicotinic acetylcholine receptors on macrophages^16^. Splenic neural signaling can also enhance T cell-dependent humoral responses^2^ and modulate antiviral immunity. During LCMV infection, catecholaminergic signaling has been associated with CD8+ T-cell exhaustion^4^, whereas during MCMV infection, β2-adrenergic signaling regulates adaptive natural killer-cell expansion and memory^3^. Thus, the consequences of splenic sympathetic activity depend on immune context and cellular target.

Splenic sympathetic innervation is also remodeled with age and disease. Catecholaminergic fiber density decreases with age in rodents and humans^17,18^, redistributes toward hilar and red-pulp regions in a mouse model of rheumatoid arthritis^19^, and is reduced in spleens from patients who died of sepsis^20^. These observations have motivated bioelectronic approaches to immune-mediated inflammatory diseases. Vagus nerve stimulation has improved clinical measures in studies of Crohn’s disease and rheumatoid arthritis^21,22^, and preclinical work has explored its potential in multiple sclerosis^23^. Recent clinical studies have further evaluated vagus nerve stimulation for rheumatoid arthritis^24^. Defining the underlying anatomy is therefore important for interpreting and refining neuromodulatory interventions.

Neural regulation of splenic immunity is unlikely to be confined to a single peripheral reflex. The inflammatory reflex is proposed to recruit a vagal-to-sympathetic pathway that engages splenic T cells through the CMG^16^. Central circuits can also influence splenic function, but the canonical sympathetic routes to the spleen and neighboring pancreas, and the descending pathways that reach CMG-projecting spinal neurons, remain poorly characterized.

In this study, we assembled a multiscale anatomical map of the descending sympathetic innervation of mouse spleen and pancreas. We visualized catecholaminergic innervation of both organs, mapped their postganglionic neurons across the full CMG complex, and generated a single-nucleus transcriptomic reference for CMG cell types. We then quantified the spinal distribution of CMG-projecting sympathetic preganglionic neurons and used glycoprotein-complemented rabies tracing to identify candidate upstream populations. The data reveal some segregation between spleen- and pancreas-projecting CMG neurons despite the proximity of the organs, identified CMG-projecting neurons across three established sympathetic regions of the thoracic spinal cord, and mapped the brainstem, hypothalamic, and cortical regions innervating these spinal circuits. Interestingly, rabies tracing revealed that the motor cortex directly innervated pre-ganglionic neurons. Complementary anterograde tracing showed motor-cortical axons and vGlut1-positive boutons in close apposition to CMG-projecting spinal neurons. These findings provide an anatomical framework of how descending circuits innervate the CMG and spleen and pancreas and suggest that descending circuits from motor cortex to the spinal cord can have effector functions beyond movement.

## RESULTS

### The spleen and pancreas can be innervated by distinct populations of CMG neurons

In mice, the splenic nerve extends from the CMG to the spleen while traversing the pancreas. We visualized catecholaminergic innervation in the spleen and pancreas by combining iDISCO tissue clearing with tyrosine hydroxylase immunolabeling^25^ (**Figure 1A; Supplementary Video 1**). Although both organs receive input from the CMG, their innervation patterns and densities differed markedly. Three-dimensional reconstructions showed that the splenic nerve contains axons continuing toward the spleen as well as branches entering the pancreas, raising the possibility that some postganglionic neurons might innervate both organs.

**Figure 1.**
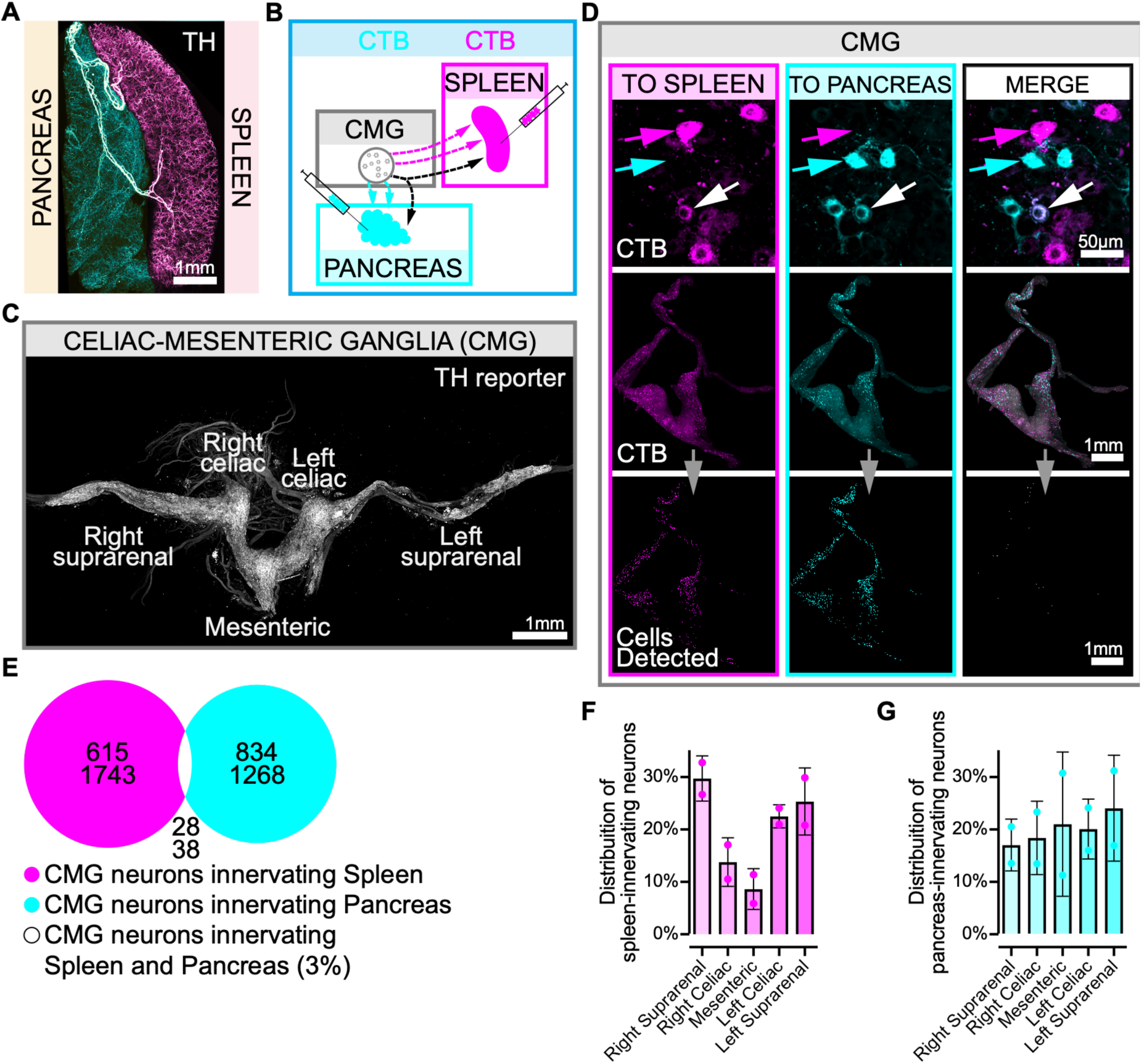
The spleen and pancreas are richly innervated by sympathetic neurons residing in the Celiac-Mesenteric Ganglia (CMG). (A) Maximum Intensity Projection of whole mount adult cleared spleen and pancreas stained with Tyrosine Hydroxylase (TH), and pseudocolored for each organ. (B) Experimental design. Cholera Toxin subunit B conjugated with Alexa Fluor 555 and 647 was injected in the spleen and pancreas to retrogradely label ganglionic neurons. (C) Whole mount of a CMG of a TH^Cre^;tdTomato^flox^ is shown, identifying the different regions of these ganglia. (D) Retrogradely labeled neurons in the CMG reveal that the vast majority of neurons project to either spleen or pancreas. (E) Venn diagram illustrating that 3.37%±1.68% of splenic neurons and 3.18%±0.25% of pancreatic neurons project to both organs. Quantifications were performed by imaging whole CMGs (middle row in D) and automatically detecting cells (bottom row in D). The halo-ring area corresponds to the standard deviation for each circle. (F) Distribution of neurons innervating the spleen in the different topographic regions of the CMG. (G) Distribution of neurons innervating the pancreas in the different topographic regions of the CMG. N=2 mice.

To identify postganglionic neurons innervating the spleen and pancreas, we injected cholera toxin subunit B (CTB) conjugated to different fluorophores into the two organs (**Figure 1B; Extended Data Figure 1A,B**). In mice, the superior mesenteric and celiac ganglia form a continuous CMG complex, which in whole-mount preparations from TH^Cre^;tdTomato^flox^ mice also appeared continuous with the suprarenal ganglion (**Figure 1C; Supplementary Video 2**). Most retrogradely labeled CMG neurons contained tracer from only one organ (**Figure 1D,E; Supplementary Video 2**). Across the analyzed animals, 3.4% ± 1.7% of spleen-labeled neurons and 3.18% ± 0.25% of pancreas-labeled neurons were double labeled (**Figure 1E**). These measurements indicate substantial organ-level segregation within the CMG. Because the analysis included only two animals and the total number of potentially labelable CMG neurons was not independently established, the data do not determine whether the small double-labeled fraction exceeds chance overlap.

We next divided the CMG into five anatomically defined regions (**Figure 1F,G**). Spleen-projecting neurons were enriched in suprarenal regions across animals (**Figure 1F**), consistent with observations in rats^26^. By contrast, pancreas-projecting neurons were distributed more broadly, with greater inter-animal variability (**Figure 1G**).

Consistent with previous reports, CTB-labeled neurons were also present in the sensory nodose-jugular complex (**Extended Data Figure 1C; Supplementary Video 3**)^27,5^. Fewer vagal sensory neurons were labeled from the spleen than from the pancreas.

### Transcriptomic characterization of CMG neuron types

To identify neuronal cell types residing in the CMG, and identify the types that project to the spleen, we collected CMG nuclei and performed single-nucleus RNA-sequencing or single-nucleus multiome. We focused on single-nucleus isolation as it is less sensitive to selective cell type depletion due to differences in cell survival (). We collected the snMultiome (10x Genomics) from CMG dissections to obtain broad cell type composition of CMG and surrounding tissue. To enrich for CMG neurons, we performed Smart-seq v4 (SSv4, Takara) snRNA-seq on nuclei obtained by fluorescence activated nucleus sorting (FANS) (**Supplementary Video 4**). The GFP+ nuclei were FANS isolated from the CMG microdissected from a Th^Cre^;Sun1-sfGFP double transgenic animal. Tyrosine hydroxylase (Th) is known to be expressed in neurons that synthesize catecholamines and most sympathetic neurons are catecholaminergic^28^.

Starting from 81,563 single nuclei (81,104 10x snMultiome and 458 SSv4), we integrated the two snRNA-seq datasets and filtered the combined data to a set of putative noradrenergic neurons (314 nuclei, or 0.4% retained; 146 10x snMultiome and 168 SSv4) (**Figure 2A; Extended Figure 2A**). In this combined neuronal dataset, we identified three clusters (**Figure 2B**), each containing nuclei from both technology platforms (**Extended Figure 2G**). Cluster 1_CMG_Rxfp1 corresponds to the previously identified Rxfp1+ neuron types^29^. Clusters 2_CMG_Robo2 and 3_CMG_Opcml correspond to subclusters of the previously identified Shox2^+^ ^29,30^ (**Extended Figure 2D,E**).

**Figure 2.**
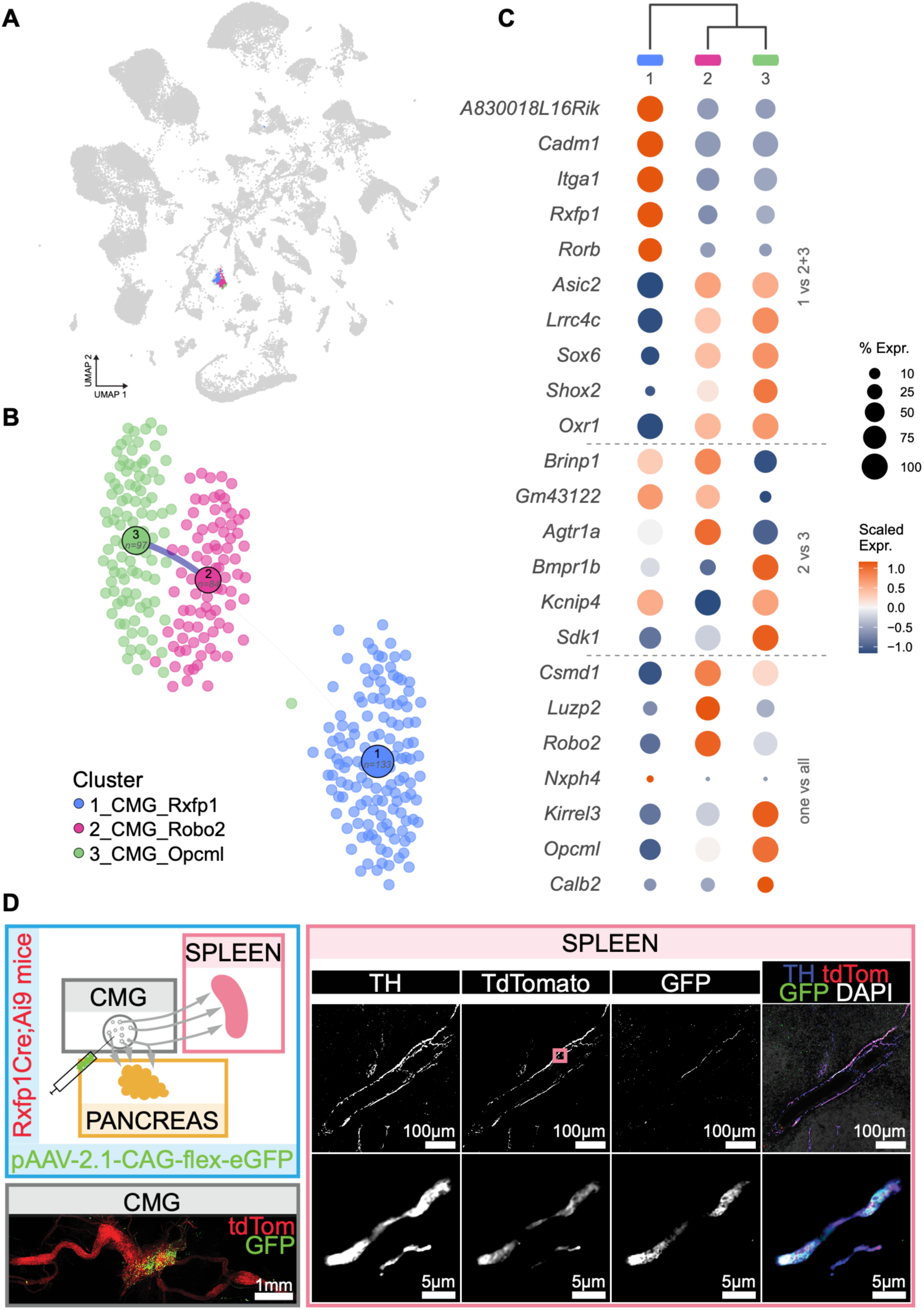
Transcriptional identity and chromatin landscape of noradrenergic CMG neurons, including spleen projections of Rxfp1^+^ neurons. (A) UMAP of 79,704 isolated single nuclei from CMG, with the retained neurons shown in color. (B) Cluster connectivity graph of the retained 314 neuronal nuclei, colored by cluster. Node size is the number of nuclei per cluster and edge weight is based on the number of cross cluster k-nearest-neighbor links, computed on the UMAP embedding (k=15). (C) Cluster marker dot plot, grouped by dendrogram contrasts. Dot size is the percent of nuclei expressing the gene and color is the scaled mean expression. (D) Spleen innervation by Rxfp1-expressing neurons. AAV-2.1-CAG-flex-eGFP was injected into the left CMG of Rxfp1-Cre;tdTomato^flox^ mice. Nearly all TH^+^ axons in the spleen expressed tdTomato, and some expressed GFP.

To determine which neurons project to the spleen, we identified an existing transgenic Cre line that corresponds to one of the marker genes identified in our transcriptomic clusters. We expect Rxfp1-P2A-Cre would label cluster 1_CMG_Rxfp1^31^. To this end, we injected the CMG of Rxfp1-Cre;tdTomato^flox^ mice with AAV expressing a reporter. Rxfp1-Cre showed projections to the spleen (**Figure 2E**), suggesting that cells from cluster 1_CMG_Rxfp1 project to the spleen, as well as other previously identified targets^29^.

### CMG-projecting sympathetic preganglionic neurons occupy three thoracic spinal regions

To map sympathetic preganglionic neurons (SPNs) that project to the CMG, we injected CTB into the ganglion complex (**Figure 3A,B**), reconstructed thoracic spinal segments with SpinalJ, and identified and quantified retrogradely labeled neurons^32^ (**Figure 3C,D**). Labeled neurons were found in the intermediolateral cell column (IML), the intercalated nucleus (ICl), and the spinal nucleus (10Sp) (**Figure 3C,D; Extended Figure 3**). This distribution extends beyond the IML pattern suggested by earlier polysynaptic tracing from the spleen^33^ and agrees with recent mapping of sympathetic spinal populations^34^. A complementary experiment using non-pseudotyped, glycoprotein-deleted N2c rabies virus produced a similar pattern (**Extended Figure 4**). In addition to spinal sympathetic inputs, CTB labeling was detected in the dorsal motor nucleus of the vagus (10N) (**Extended Figure 5**), consistent with previously described vagal projections to the CMG^35^.

**Figure 3.**
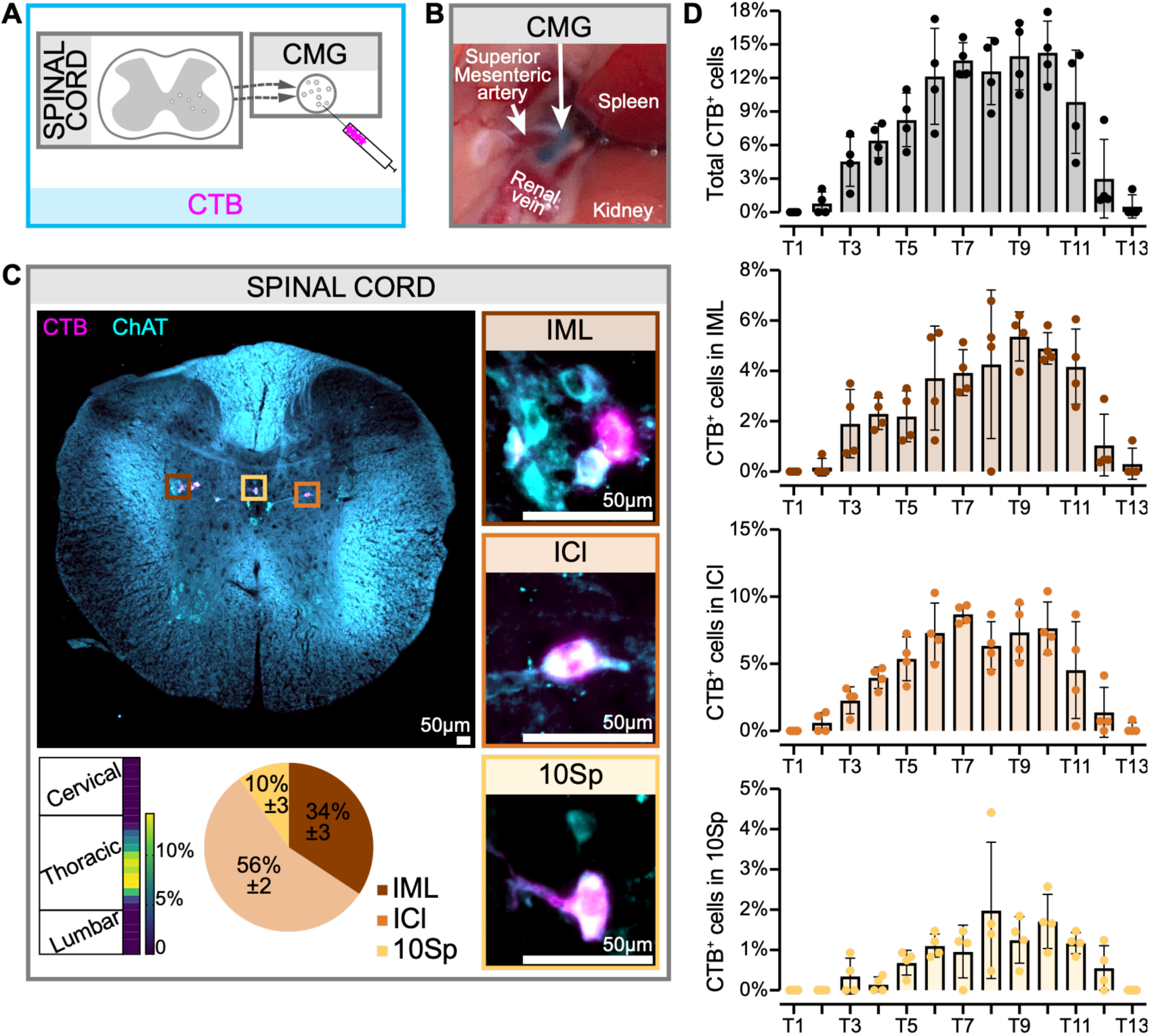
The Celiac-Mesenteric Ganglia (CMG) is innervated by neurons residing in known preganglionic sites between T3 and T12 of the thoracic spinal cord. (A) Experimental design. Cholera Toxin subunit B was injected in the CMG to retroactively label preganglionic neurons in the spinal cord. (B) The CMG is visibly blue after injection of Alexa Fluor-647 conjugated CTB. (C) Neurons are found only in the thoracic segments of the spinal cord. Middle, thoracic cord section containing CTB^+^ preganglionic ChAT^+^ neurons in the intermediolateral nucleus (IML), intercalated nucleus (ICl) and lamina X (10Sp). Bottom left, heatmap representing the distribution of CTB^+^ neurons throughout the spinal cord. Bottom center, pie chart representing the distribution of CTB^+^ neurons in different spinal cord regions. (D) Total, IML, ICl, and 10Sp percentage of counted CTB^+^ neurons per thoracic spinal cord segment. N=4 mice.

### Neurons across various brain regions directly innervate CMG-projecting preganglionic neurons

To identify brain regions containing neurons upstream of CMG-projecting spinal populations, we used a glycoprotein-complemented rabies tracing strategy. We injected a Cre-dependent helper AAV expressing N2cG into thoracic spinal segments T3-T11 of ChAT-Cre mice, guided by the distribution of CMG-projecting neurons described above, and subsequently injected non-pseudotyped rabies virus into the CMG (**Figure 4A**). Rabies virus taken up by axon terminals in the CMG could reach spinal neurons retrogradely; in ChAT-positive neurons co-transduced with the helper AAV, glycoprotein complementation permitted further retrograde spread. Whole-brain reconstruction with BrainJ identified labeled neurons in the brainstem, hypothalamus, and cortex (**Figure 4A; Supplementary Video 5**). Most labeled cells were in the medulla (60% ± 9%), followed by the cortex (21% ± 4%), pons (13% ± 6%), midbrain (3% ± 1%), and hypothalamus (0.8% ± 0.6%) (**Figure 4B; Extended Figure 6**). Within the brainstem, labeled neurons occurred mostly in the reticular formation. The majority of neurons in the hypothalamus occurred in the paraventricular hypothalamic nucleus (PVH). Within the cortex, labeled neurons occurred in somatosensory, motor, and frontal regions, and some labeled corticospinal neurons had striatal collaterals (**Figure 4B**).

**Figure 4.**
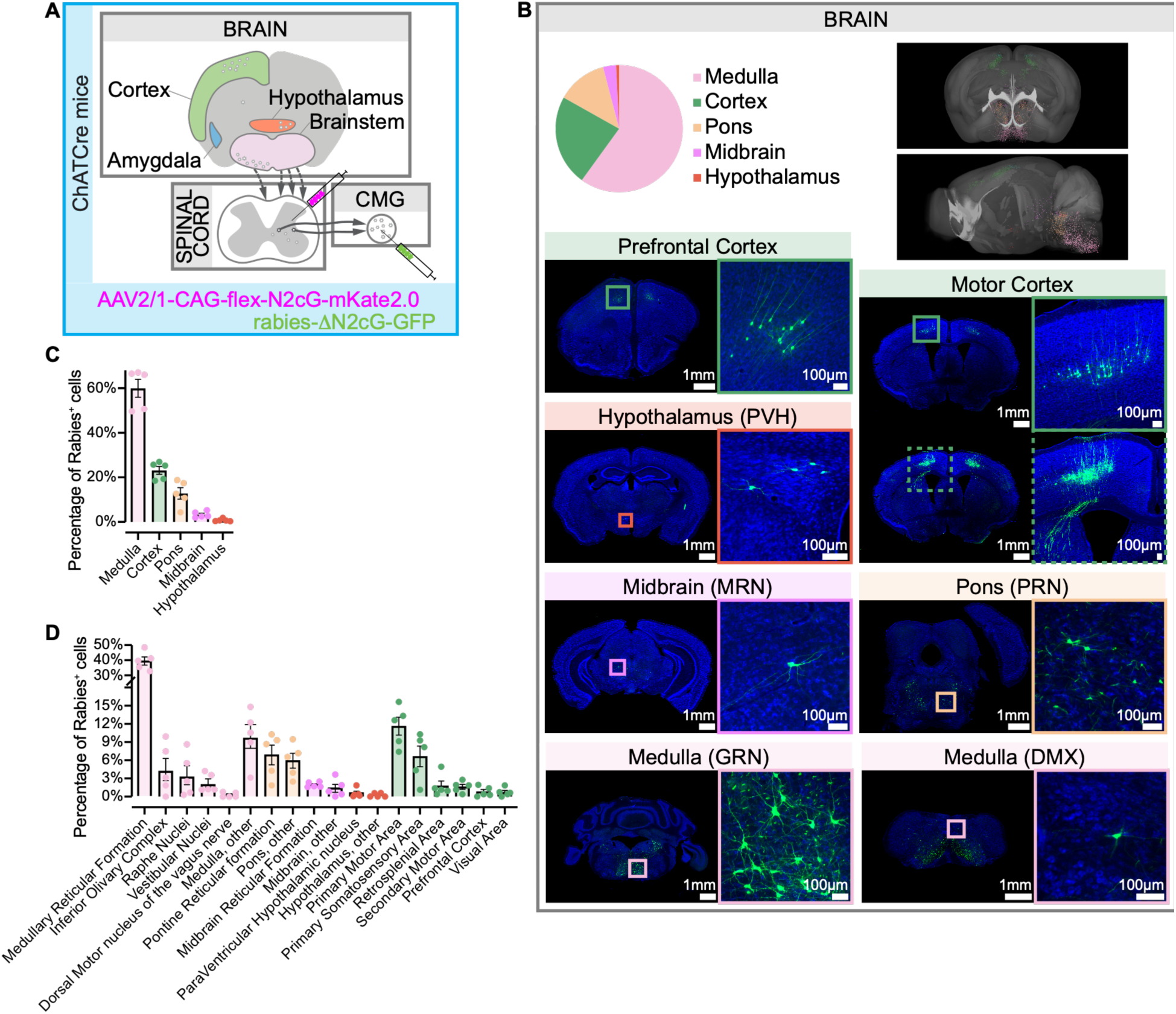
The Celiac-Mesenteric Ganglia (CMG) preganglionic spinal neurons are innervated by neurons residing in brainstem, hypothalamus and cortex. (A) Experimental design. A Cre-dependent AAV expressing rabies G protein was injected into the spinal cord of Chat-Cre mice. 6 weeks later, G-deficient rabies expressing GFP was injected into the CMG. (B) Top left, pie chart representing the distribution of rabies^+^ cells in the brain, 10 days after rabies injection. Top right, example of a representative brain revealing the localization of rabies^+^ cells within the brain. Bottom, representative images of rabies^+^ cells in different brain regions. (C) Quantification of rabies^+^ neurons in the 5 brain structures where any rabies^+^ neuron was found. (D) Quantification of rabies^+^ neurons in the brain regions identified. Medullary Reticular Formation is GRN, MARN, PGRNd, PGRNl, IRN, LRNm, MDRNv, PARN and MDRNd. Raphe Nuclei is RO, RM and RPA. Vestibular Nuclei is MV, LAV, SPIV and SUV. Pontine Reticular Formation is PRNc, PRNr and TRN. Midbrain Reticular Formation is MRN and RN. N=5 mice.

### Motor cortex neurons synapse in the brain in close apposition to CMG-projecting preganglionic neurons

Given the surprising results of sensorimotor cortex innervating spinal neurons that project to CMG, we wanted to check for the possibility that our rabies strategy allowed the rabies to jump multiple synapses between ChAT^+^ cells in the spinal cord before reaching the brain. We therefore perfect experiments seeking orthogonal anatomical evidence for this observation. To do so, we injected AAV expressing a reporter into the motor cortex of WT mice. We then injected CTB into the CMG of the same mice (**Figure 5A**). We localized CTB^+^ CMG-projecting SPNs in proximity with the synaptic market vGlut1 of motor cortex corticospinal axons (**Figure 5B**). This anatomical proximity between motor corticospinal synapses and CMG-projecting SPNs were found in all three regions of the spinal cord – 10Sp, ICl and IML (**Figure 5B**).

**Figure 5.**
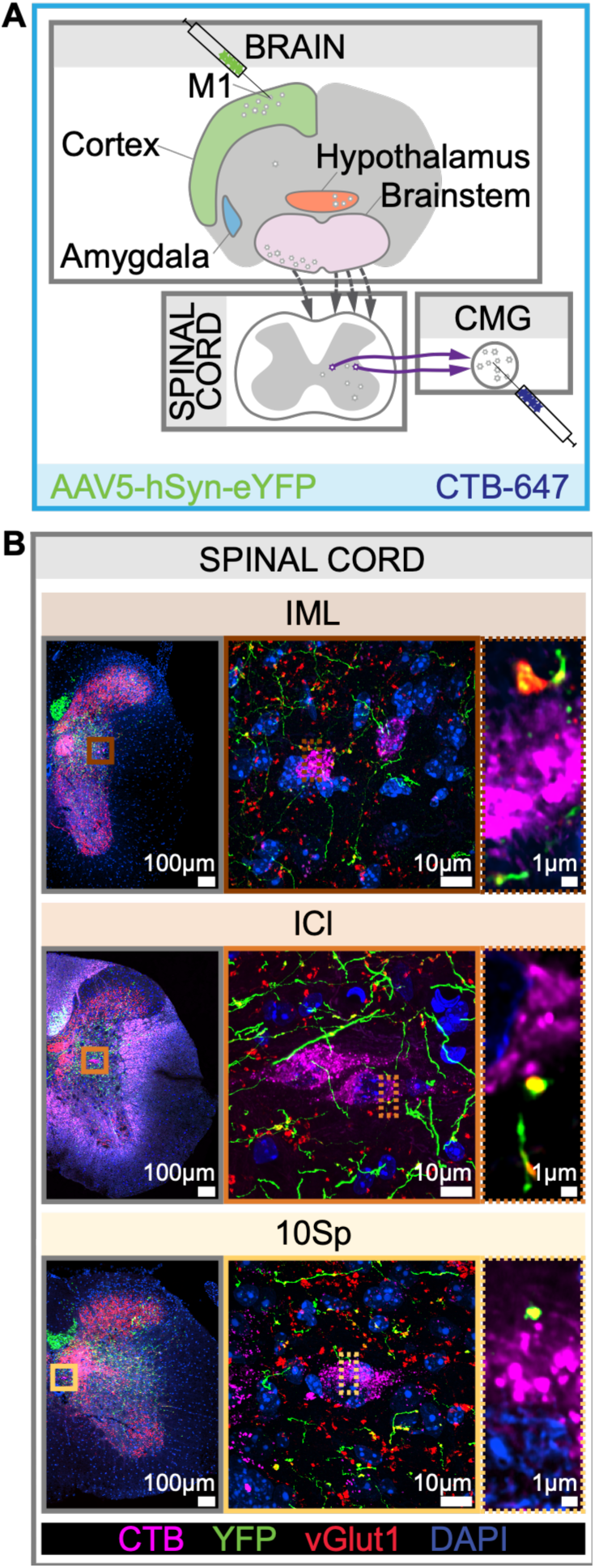
Motor cortex neurons synapse in the brain in close apposition to CMG-projecting preganglionic neurons. (A) Experimental design. AAV expressing YFP was injected directly into primary motor cortex (M1). Three weeks later, CTB was injected into the CMG. (B) Spinal cord sections of these mice 7 days after CTB injection showing apposition between synaptic sites of corticospinal neurons projecting from motor cortex and CMG-projecting preganglionic neurons in IML, ICl and 10Sp.

## DISCUSSION

Sympathetic innervation of the spleen modulates vascular and immune functions^36^, yet the anatomical organization of the pathway from the brain to splenic postganglionic neurons has remained fragmented across studies. Here, we combine retrograde tracing, single-nucleus profiling of the CMG, quantitative spinal mapping, and transsynaptic tracing to provide an integrated framework spanning the spleen and pancreas, CMG, thoracic spinal cord, and brain. Three findings are particularly informative: spleen-and pancreas-projecting neurons are largely segregated within the CMG; CMG-projecting SPNs occupy 10Sp, ICl, and IML; and candidate descending inputs arise from distributed brainstem, hypothalamic and cortical regions.

Analysis of the complete CMG preparation showed that only 3.4% ± 1.7% of spleen-labeled neurons and 3.18% ± 0.25% of pancreas-labeled neurons contained both tracers. The predominant pattern is therefore organ-selective projection, consistent with recent findings in other abdominal sympathetic pathways^29^. The small double-labeled fraction could reflect genuine axonal branching, tracer spread, incomplete sampling, or chance overlap. Because the present dataset does not provide an independent estimate of all neurons available for labeling and includes two animals, it cannot distinguish among these possibilities. If validated with orthogonal approaches, true dual-projecting neurons would be candidates for coordinating immune and metabolic functions. Analogous dichotomizing sensory neurons have been proposed to contribute to cross-organ signaling between the colon and bladder^37^.

Most spleen-projecting neurons were located in splanchnic and suprarenal regions of the CMG complex (**Figure 1**), consistent with the preferential localization of spleen-projecting neurons in rat suprarenal ganglia^26^. A previous mouse study localized spleen-and pancreas-projecting neurons primarily to celiac portions of the CMG but did not include the splanchnic or suprarenal components^29^. Our whole-mount preparation included the splanchnic nerve and plexus, suprarenal ganglia, and aortico-renal plexus^38^, revealing a broader distribution. Comparative anatomical work identifies related celiac structures in fish and amphibians^39–41^, whereas the human celiac ganglia are more discretely defined and the superior mesenteric ganglion is less prominent^42^. Although homologies among these structures require caution, the conserved association of celiac ganglia with major abdominal vessels suggests an evolutionarily retained organizational scaffold for visceral sympathetic output.

In line with previous data, the CMG cluster that contains neurons projecting to the spleen, Rxfp1^+^, expresses the α7 nicotinic acetylcholine receptor (**Figure 2B**; **Extended Data Figure 2**)^43^. CMG-projecting SPNs occupied 10Sp, ICl, and IML, in agreement with recent anatomical mapping^34^. Earlier polysynaptic tracing from the spleen emphasized the IML^33^, a difference that may reflect viral tropism, route of administration, survival time, or the broader set of CMG targets labeled here. Comparative studies also suggest that the distribution of sympathetic preganglionic neurons across homologous spinal regions varies among vertebrates: amphibian celiac pathways arise predominantly from IML neurons, whereas broader surveys identify neurons in IML and IC in amphibians and in central gray regions in fish^44^.

Polysynaptic pseudorabies-virus studies of the spleen, pancreas, colon, and celiac ganglia have labeled neurons in the brainstem and hypothalamus, and some studies have also reported prefrontal cortical labeling^33,45–47^. Rabies-based tracing from the adrenal gland, stomach, and kidney has identified cortical populations, including layer 5 motor-cortical neurons^38,47–49^. Differences among reports may reflect organ specificity as well as distinct viral tropisms and tracing kinetics. Our dataset identifies the medulla as the largest source of labeled brain neurons, followed by additional populations in the pons, midbrain, hypothalamus, and cortex. These distributed populations are anatomically positioned to integrate homeostatic, arousal, sensorimotor, and executive signals before sympathetic output reaches abdominal organs. Determining which populations connect specifically to spleen-versus pancreas-projecting pathways will require organ-restricted intersectional tracing.

Motor-cortical axons and vGlut1-positive boutons were closely apposed to CTB-labeled CMG-projecting neurons in 10Sp, ICl, and IML. Together with the rabies-labeling pattern, this observation supports a candidate route through which motor-related cortical activity could access sympathetic spinal circuits. However, neither light-microscopic apposition nor the present viral strategy proves direct, functional synapses onto CMG-projecting SPNs. Electron microscopy, ex vivo circuit mapping, or projection-specific physiological manipulation will be needed to establish connectivity and function. The current results therefore broaden the anatomical framework for motor-cortical output beyond somatic motor control while providing testable hypotheses about cortical regulation of visceral physiology, including immune function.

## METHODS

### Experimental model and animals

All experiments and procedures were performed according to National Institutes of Health (NIH) guidelines and approved by the Institutional Animal Care and Use Committee of Columbia University. Adult mice of both sexes, aged 1.5–6 months, were used for all experiments. The strains used were: C57BL/6J (Jackson Laboratories, 000664), B6.129S-Chat^tm1(cre)Lowl^/MwarJ (Jackson Laboratories, 031661; identified in this paper as Chat^Cre^), B6.FVB(Cg)-Tg(Th-cre)FI172Gsat/Mmucd (MMRRC, 031029-UCD; identified in this paper as TH^Cre^), B6.129-Gt(ROSA)26Sor^tm5.1(CAG-Sun1/sfGFP)Nat^/MmbeJ (Jackson Laboratories, 030952; identified in this paper as CAG-Sun1/sfGFP), B6.Cg-Rxfp1^em1(cre)Ngai^/TasicJ (Jackson Laboratories, 037336), and B6;129S6-Gt(ROSA)26Sor^tm9(CAG-tdTomato)Hze^/J (Jackson Laboratories, 007905; identified in this paper as tdTomato^flox^). All Cre lines used were heterozygotes crossed to C57BL/6J mice. All mice used in experiments were individually or pair housed under a 12-h light/dark cycle.

### Stereotaxic viral injections

Buprenorphine SR (1 mg/kg) was administered subcutaneously on the day of the surgery. Mice were anesthetized with isoflurane and placed in a spinal stereotaxic frame (Kopf). Eye cream was applied, the fur was shaved on the surgical site, and the skin was cleaned alternating ethanol and Iodine (Βetadine) swabs. Bupivacaine (2 mg/kg) was administered subcutaneously on the surgical site. All injections were performed using glass pipettes (beveled using a Kimwipe for all injections but blunted for brain) mounted to a Nanoject (Nanoject III, Drummond).

For pancreas, spleen, and CMG surgeries, the mice were placed on a prone position. Lactated Ringer’s solution (Covetrus 25211-8) was used to keep the abdomen humidified throughout all abdominal surgeries. For spleen and pancreas surgeries, a left lateral laparotomy was made over the injection sites. The organs were held in place using a retractor modified by inverting its angle (F.S.T. 17004-03) fixed to the spinal stereotaxic frame using a spinal clamp. Appropriate pancreas injection was confirmed by confirming proper CTB spread within the tissue (and not in the folds) upon the first injection pulse. Care was made to ensure there was no efflux of virus by stabilizing the skull and waiting 2 min after penetration before and after injecting. Each site was injected with 500nL of virus; 8 injections distributed throught the spleen for a total of 4µL, 4 injections in different sites of the pancreas for a total of 2µL.

For CMG injections, a midline laparotomy was performed, and the back of the abdominal cavity was accessed by gently pulling all abdominal organs to the side of the cavity using wire retractors. The CMG was placed vertically for appropriate injections and gently cleaned using mini cotton tips (Puritan 25-826 5WC) prior injections and after each injection.

For spinal cord and brain surgeries, the mice were placed on a supine position. For spinal cord surgeries, a midline incision was made to expose the thoracic spinal column. The spinalis muscles were retracted to gain access to the T2 thoracic process which was clamped to stabilize the spinal cord using a crocodile clamp fixed to the spinal stereotaxic frame using a spinal clamp. The tail was gently stretched with another spinal clamp to separate the vertebrae. The ligaments between the laminae were removed using fine forceps. A needle was used to sever the meninges, exposing the spinal cord. 250nL of 9.83×1012vg/mL AAV2/1-CAG-Flex-N2c-G-mKate2.0 (Janelia) of 4.87×10^12^vg/mL AAV2/2-CAGGS-FLEX-Rev-mKate-T2A-CVS-N2cG (UNC) were injected in 6 sites of the spinal cord: (1) caudal between T3 and T4 intervertebral disks, (2) rostral and (3) caudal between T4 and T5 intervertebral disks, (4) between T5 and T6 intervertebral disks, (5) between T6 and T7 intervertebral disks, and (6) between T7 and T8 intervertebral disks to target T5 to T10 spinal cord segments. All injections were performed 250µm lateral to the midline, 550 µL below the cord surface.

For brain surgeries, a midline incision was made to expose the skull, and a craniotomy was made over the injection site. To label M1 corticospinal neurons, 30nL of 5.6×10^12^vg/nL AAV5-hSyn-eYFP (UNC) was injected in 4 sites of motor cortex, 600µm below the pial surface: 950µm anterior, 1mm lateral to Bregma; 350µm anterior, 0.9mm lateral to Bregma; 0.13 posterior, 1.14mm lateral to Bregma; and 610µm posterior, 1.4mm lateral to Bregma. Care was made to ensure there was no efflux of virus by stabilizing the skull and waiting 10 min after penetration before injecting. The abdominal muscle was sutured using PGA absorbable sutures (5/0) and the skin sutured with Nylon sutures (5/0).

### Histology and lightsheet imaging

Animals were deeply anesthetized with Isoflurane and transcardially perfused with 0.1M PBS 10U/mL Heparin followed by ice-cold 4% Paraformaldehyde (PFA, Electron Microscopy Sciences, 19208) in phosphate buffer (Sigma-Aldrich, P3619). Spleen and tail of the pancreas were extracted as a whole and post-fixed overnight in 4% PFA at 4°C. iDisco protocol was followed immediately after as described in https://idisco.info/idisco-protocol/ (Renier 2014). Permeabilization and blocking were done in 2 day long incubations. Primary antibody rabbit anti-TH (Millipore AB152, 1:1,000) was incubated for 5 days. Secondary antibody donkey anti-rabbit 647 (Invitrogen A31573, 1:1,000) was incubated for 5 days. Data was processed using Imaris.

### Histology and confocal imaging

Animals were deeply anesthetized with Isoflurane and transcardially perfused with 0.1M PBS 10U/mL Heparin followed by ice-cold 4% Paraformaldehyde (PFA, Electron Microscopy Sciences 19208) in phosphate buffer (Sigma-Aldrich, P3619). Spleen, pancreas, undissected ganglia, and brains were extracted and post-fixed overnight in 4% PFA at 4°C and stored in PBS 0.05% Thimerosal at 4°C until further processing. Spinal cords were collected and post-fixed for 2 hours in 4% PFA at 4°C and processed for freezing. All tissues to be cryopreserved were dehydrated in 30% sucrose solution at 4°C until sunk, transferred to Optimum Cutting Temperature Compound (Sakura Finetek, 4583) and frozen in a slush of ethanol and crushed dry ice. Spinal cords were prepared as described in Fiederling 2021. The tissue was rinsed several times in PBS, then permeabilized for 1 hour in PBS containing 0.2% Triton X-100 and 6% donkey serum. Immunostaining was performed in PBS 0.2% Tween-20 3% donkey serum with primary antibodies overnight at 4°C and secondary antibodies 2 hours at room temperature. Antibodies used were rabbit anti-TH (Millipore AB152, 1:1,000), goat anti-CTB (Lise Labs 703, 1:8,000), rabbit anti-ChAT (Columbia University AB_2750952, 1:5,000), chicken anti-GFP (Aves Labs GFP-1020, 1:2,000), guinea pig anti-vGlut1 (Columbia University AB_2750941, 1:8,000), donkey anti-rabbit 647 (Invitrogen A31573, 1:1,000), donkey anti-goat 647 (Invitrogen A32849, 1:1,000), donkey anti-guinea 568 (Jackson ImmunoResearch 706-575-148, 1:800), donkey anti-chicken 488 (Jackson ImmunoResearch 703-545-155, 1:800), NeuroTrace 500/525 Green Fluorescent Nissl Stain (Invitrogen N21480).

Coronal brain sections were performed after fixation either in a vibratome (Leica VT1000 S) or in cryostat (Leica) after cryopreservation.

Ganglia were dissected as whole mounts. Ganglia and high-magnification images of spinal cord sections were acquired on a W-1 Yokogawa spinning disk confocal microscope using 20× or 100× oil immersion objectives. Ganglia images were denoised using NIS-Elements’ NIS.ai. Spinal cord sections and ganglia imaged with the 100× objective were additionally deconvolved in NIS-Elements. For quantification of retrogradely labeled neurons, whole ganglia were imaged with a 20× objective and NIS.ai was used to train a custom object segmentation model for labeled neurons. Each CMG whole mount was cropped around the ganglia, and 5–10 images per channel were selected for model training. Detected objects below a minimum volume threshold were excluded to eliminate debris and incomplete detections, and any detected objects outside the tissue boundary were manually removed. Labeled neurons were counted from the resulting segmentations for each whole mount preparation. Maximum intensity projection images were generated in NIS-Elements (**Figure 1A**). Z-stack videos were generated in FIJI (**Supplementary Videos 2 and 4**).

Venn diagrams were generated as area-proportional figures. Circle radii were scaled such that each circle’s area was proportional to the mean neuron count across both animals (radius ∝√n̅). The center-to-center distance between circles was determined iteratively so that the computed lens-shaped intersection area matched the mean number of dual-projecting neurons, using the standard formula for the area of a circular lens. Concentric rings around each circle represent the standard deviation of counts across animals (n = 2; ring area proportional to SD). Figures were generated as scalable vector graphics (SVG) in Python and finalized in Adobe Illustrator.

Topographic analysis was performed across five anatomically distinct CMG regions (right suprarenal, right celiac, mesenteric, left celiac, left suprarenal), defined by approximate anatomical landmarks. Suprarenal regions included the splanchnic plexus, suprarenal ganglia, and aortico-renal plexus. Each CMG image was cropped to isolate these five, and the same detected objects from the whole-mount segmentation were re-counted within each cropped region. To assess whether spleen-and pancreas-innervating neurons were non-uniformly distributed across CMG topographic regions, and whether their distributions differed from each other, we performed chi-square tests on counts of retrogradely labeled neurons across five defined regions (right suprarenal, right celiac, mesenteric, left celiac, left suprarenal).

### Single-nucleus RNA and ATAC profiling

#### Dissection and nuclear isolation for CMG

C57/Bl6 or TH^Cre^; CAG-Sun1/sfGFP mice were anesthetized with isoflurane and placed in a spinal stereotaxic frame (Kopf). A midline laparotomy was performed, and the back of the abdominal cavity was accessed by gently pulling all abdominal organs to the side of the cavity using wire retractors. The whole CMG was collected, including surrounding tissue, quickly dried off on a Kimwipe to remove excess blood and placed on a perforated spoon which was frozen in liquid nitrogen vapors by holding the spoon right above liquid nitrogen for 2 minutes. CMGs were placed in 0.5mL tubes (Sarstedt) containing 100µL O.C.T. Compound (Tissue-Tek, Sakura). Frozen samples were stored at -80⁰C until processing. Each CMG was processed with the RAISINs protocol with modifications as described (https://dx.doi.org/10.17504/protocols.io.8epv517xdl1b/v5).

#### Flow cytometry for SMART-seq v4

Following nuclear isolation, Th^Cre^;CAG-Sun1-sfGFP individual fluorescent^+^ nuclei were sorted into 8-well strips containing SMART-seq lysis buffer with RNAse inhibitor (0.17 U/µL; Takara Cat#ST0764), then frozen at -80⁰C to await processing. SMART-seq libraries were sequenced to a target read depth of 500,000 reads per nucleus, as previously described^50,51^. The full protocol for sample preparation and library preparation is documented at under “Documentation.”

#### 10x Genomics single nucleus multiome sequencing

Following nuclear isolation, nuclei were quantified and processed using 10x Genomics Chromium Next GEM Single Cell Multiome Kit (Cat# 1000234). We prepared 10x Genomics snMultiome samples as previously described https://dx.doi.org/10.17504/protocols.io.bp2l61mqrvqe/v3; https://dx.doi.org/10.17504/protocols.io.dm6gpwd8jlzp/v3).

#### Processing and data analysis

Single nucleus RNA-seq and ATAC-seq data were analyzed with the package scCairns (https://github.com/AllenInstitute/scCairns), which yields a fully reproducible and documented analysis trail (available upon publication). In brief, all RNA-seq data from SMART-seq v4 and snMultiome were combined, subjected to basic QC (minimum genes per cell = 500; minimum cells expressing a gene = 1), then integrated with Harmony^52^ and scVI^53^ across a range or model architectures. The best performing model was selected to move forward via scib-metrics scores^54^. The initial subset of putative neurons was selected using assumed markers of sympathetic neurons including (*Snap25*, *Isl1*, *Phox2b* and *Th*). Once the data were subset to a small number of neuronal clusters, several iterative rounds of re-integration and QC were performed to arrive at a final neuronal subset (**Extended Figure 2A**). QC characteristics included identifying cluster outliers for metrics such as gene count and mitochondrial ratio, clusters dominated by individual animals, or clusters not expressing neuronal markers. Final clustering resolution was determined by sweeping across a range of Leiden clustering resolutions and maximizing the silhouette score. Differential expression was calculated in Seurat with FindAllMarkers.

## ACKNOWLEDGEMENTS

This research was supported by the NIH Director’s Pioneer Award DP1AT011979 (to R.M.C.) and the Allen Institute and Fund for Science and Technology (FFST).

We thank Nicolas Renier for his imaging and analysis of spleen and pancreas using iDisco.

---

**Extended Data Figure 1.**
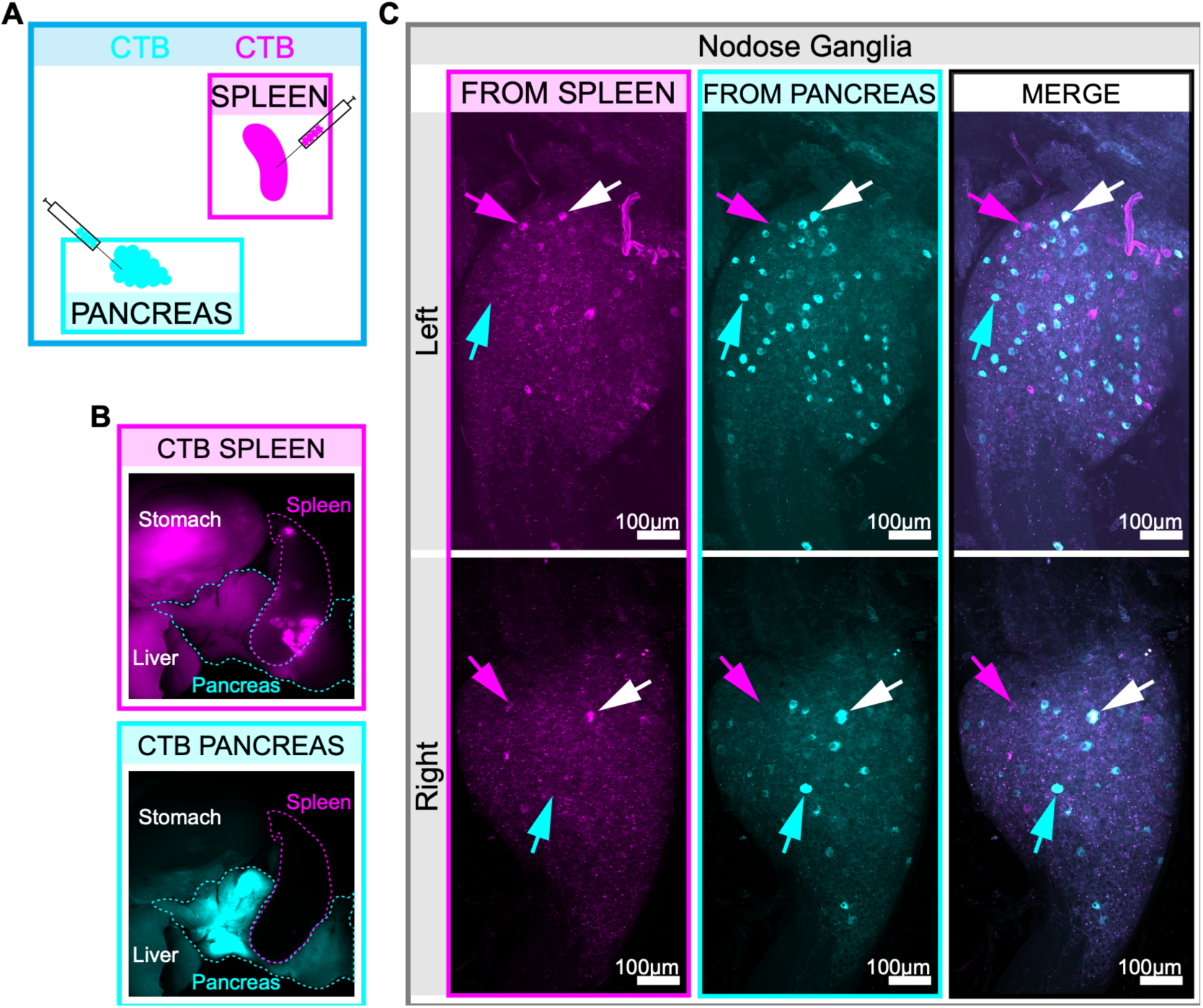
Labeling of spleen and pancreas innervating neurons. (A) Experimental design. Cholera Toxin Subunit B (CTB) conjugated with Alexa Fluor fluorophores are injected in spleen and pancreas (Figure 1). (B) Injection sites of dual colored CTB are visualized 7 days post injection. (C) CTB-labeled neurons in nodose ganglia.

**Extended Data Figure 2.**
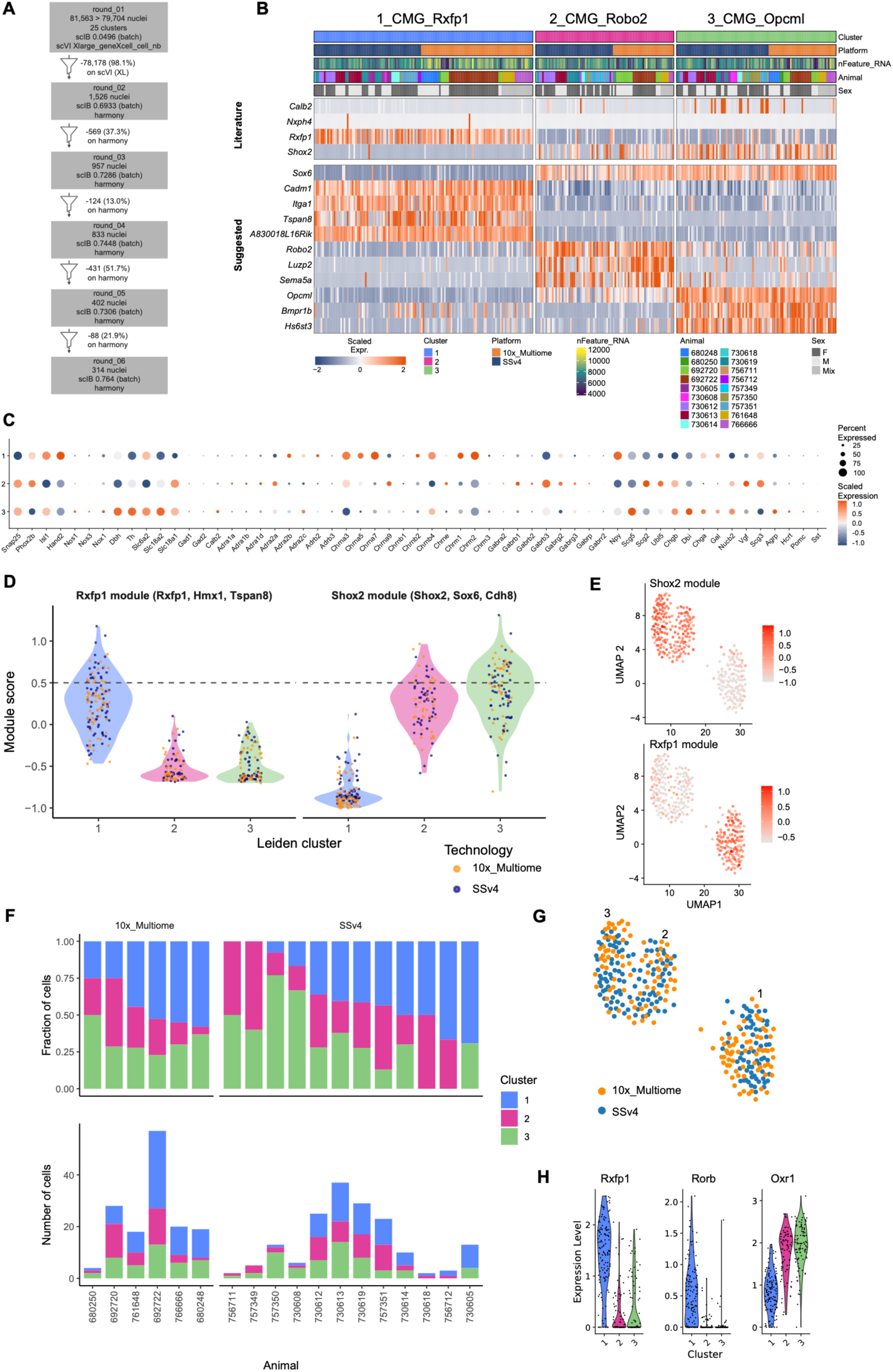
Filtering and annotation of noradrenergic CMG neurons. (A) Filtering approach from starting nucleus population to the final 314 nuclei over 6 rounds. (B) Single nucleus heatmap displaying previously defined “Literature” markers and additional suggested markers for each of the three clusters. (C) Dot plot of neuronal markers, receptors and neuropeptides by cluster. Dot size is percent of nuclei expressing and color is the scaled expression. (D) Module scores for Rxfp1-related genes and Shox2-related genes, calculated with Seurat’s AddModuleScore function. Points are colored by technology platform. Dashed line is arbitrarily set to 0.5. (E) UMAP colored by the Shox2 and Rxfp1 module scores. (F) Cluster composition per animal, with each platform separately displayed. (G) UMAP colored by technology platform. (H) Expression of Rxfp1, Rorb, and Oxr1 by cluster.

**Extended Data Figure 3.**
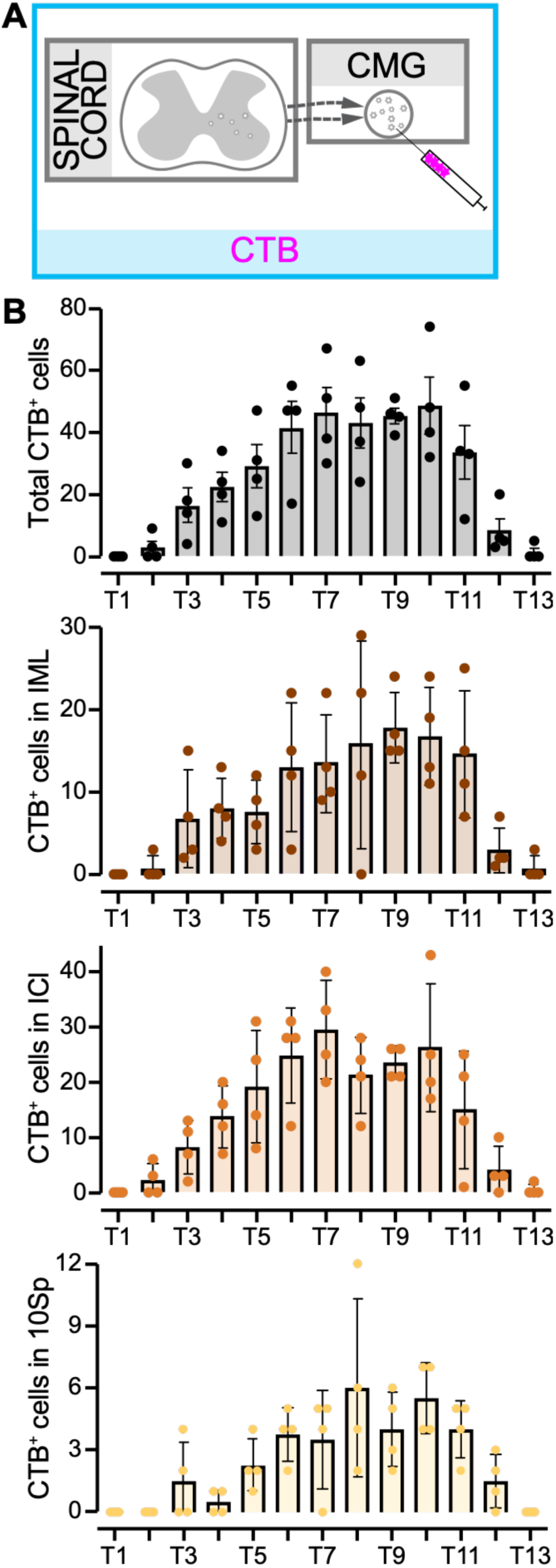
The Celiac-Mesenteric Ganglia (CMG) is innervated by neurons residing in known preganglionic sites between T3 and T12 of the thoracic spinal cord. (A) Experimental design. Cholera Toxin subunit B was injected in the CMG to retroactively label preganglionic neurons in the spinal cord. (B) Total, IML, ICl, and 10Sp number of counted CTB^+^ neurons per thoracic spinal cord segment. N=4 mice.

**Extended Data Figure 4.**
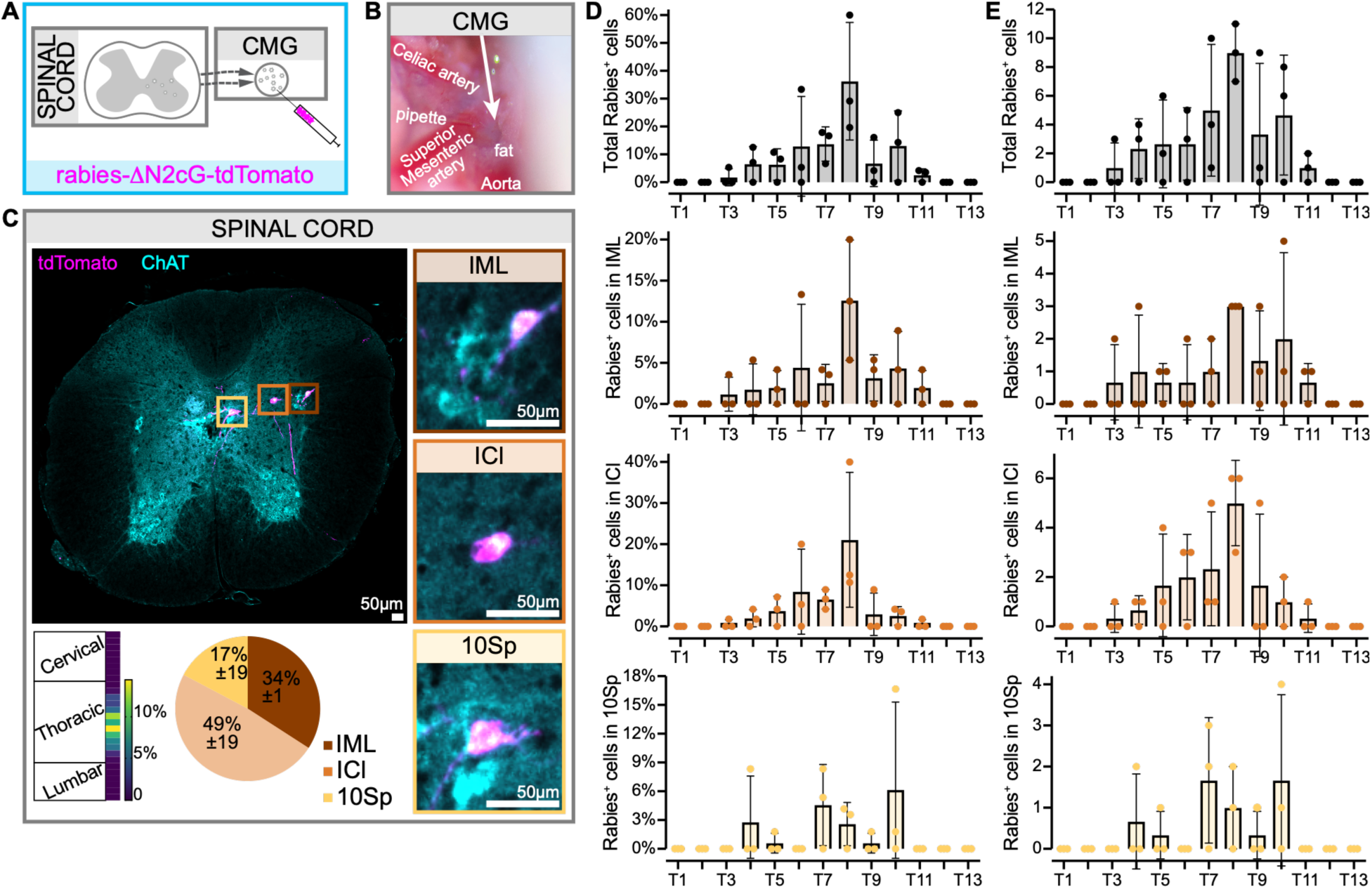
The Celiac-Mesenteric Ganglia (CMG) is innervated by neurons residing in known preganglionic sites between T3 and T12 of the thoracic spinal cord. (A) Experimental design. Nonpseudotyped rabies virus was injected in the CMG to retroactively label preganglionic neurons in the spinal cord. (B) Surgical setup showing the glass pipette in the CMG during rabies injection. (C) Neurons are found only in the thoracic segments of the spinal cord. Middle, thoracic cord section containing rabies^+^ preganglionic ChAT^+^ neurons in the intermediolateral nucleus (IML), intercalated nucleus (ICl) and lamina X (10Sp). Bottom left, heatmap representing the distribution of rabies^+^ neurons throughout the spinal cord. Bottom center, pie chart representing the distribution of rabies^+^ neurons in different spinal cord regions. (D) Total, IML, ICl, and 10Sp percentage of counted rabies^+^ neurons per thoracic spinal cord segment. (E) Total, IML, ICl, and 10Sp number of counted rabies^+^ neurons per thoracic spinal cord segment. N=3 mice.

**Extended Data Figure 5.**
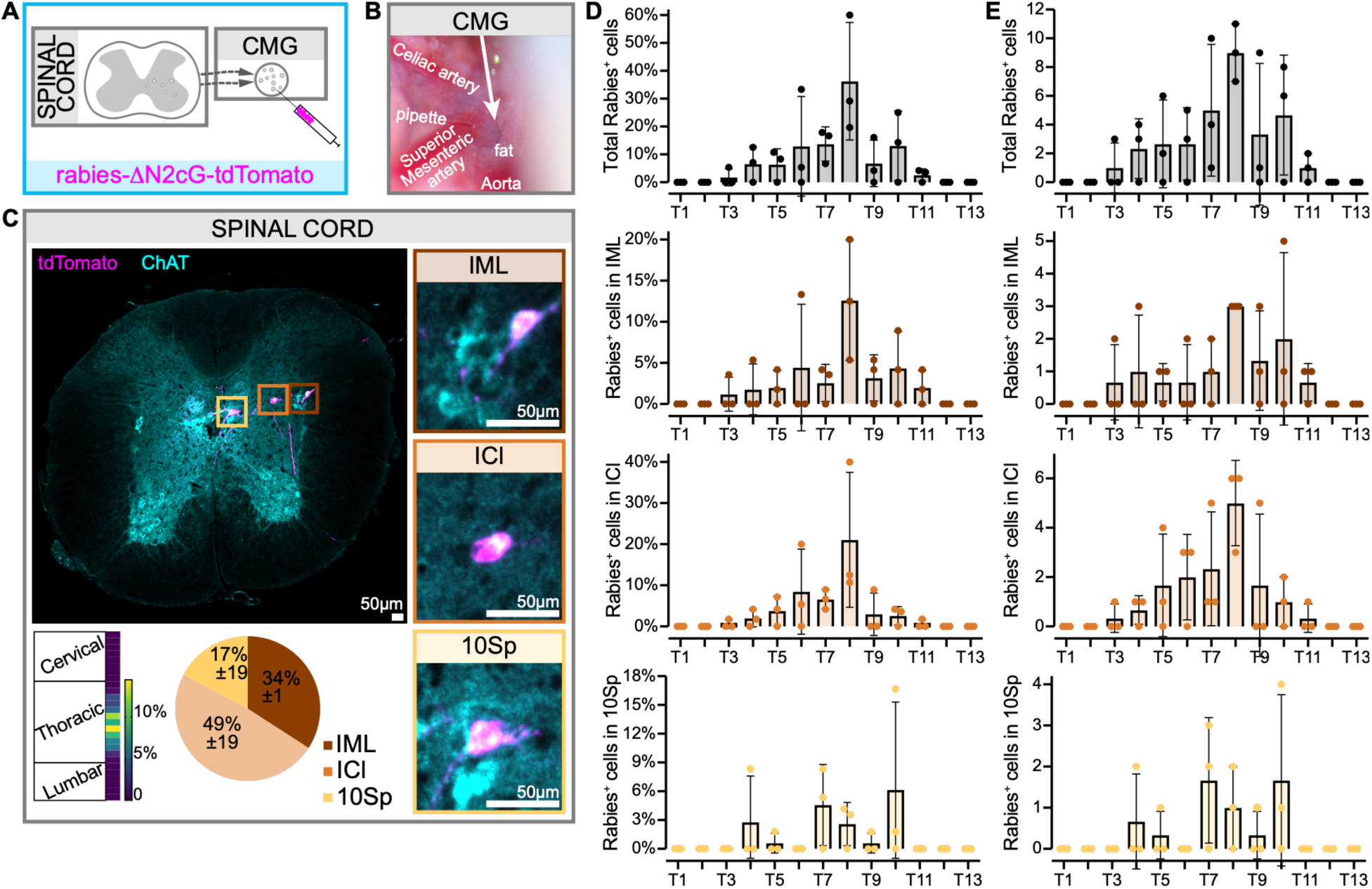
The Celiac-Mesenteric Ganglia (CMG) is innervated by neurons residing in the dorsal motor nucleus of the vagus in the brainstem (10N). (A) Experimental design. Cholera Toxin subunit B (CTB) was injected in the CMG to retroactively label preganglionic neurons in the spinal cord (Figure 3). (B) CTB^+^ neurons are found in the dorsal motor nucleus of the vagus in the brainstem. Red, antibody-labeled CTB neurons. Green, NeuroTrace 500/525.

**Extended Data Figure 6.**
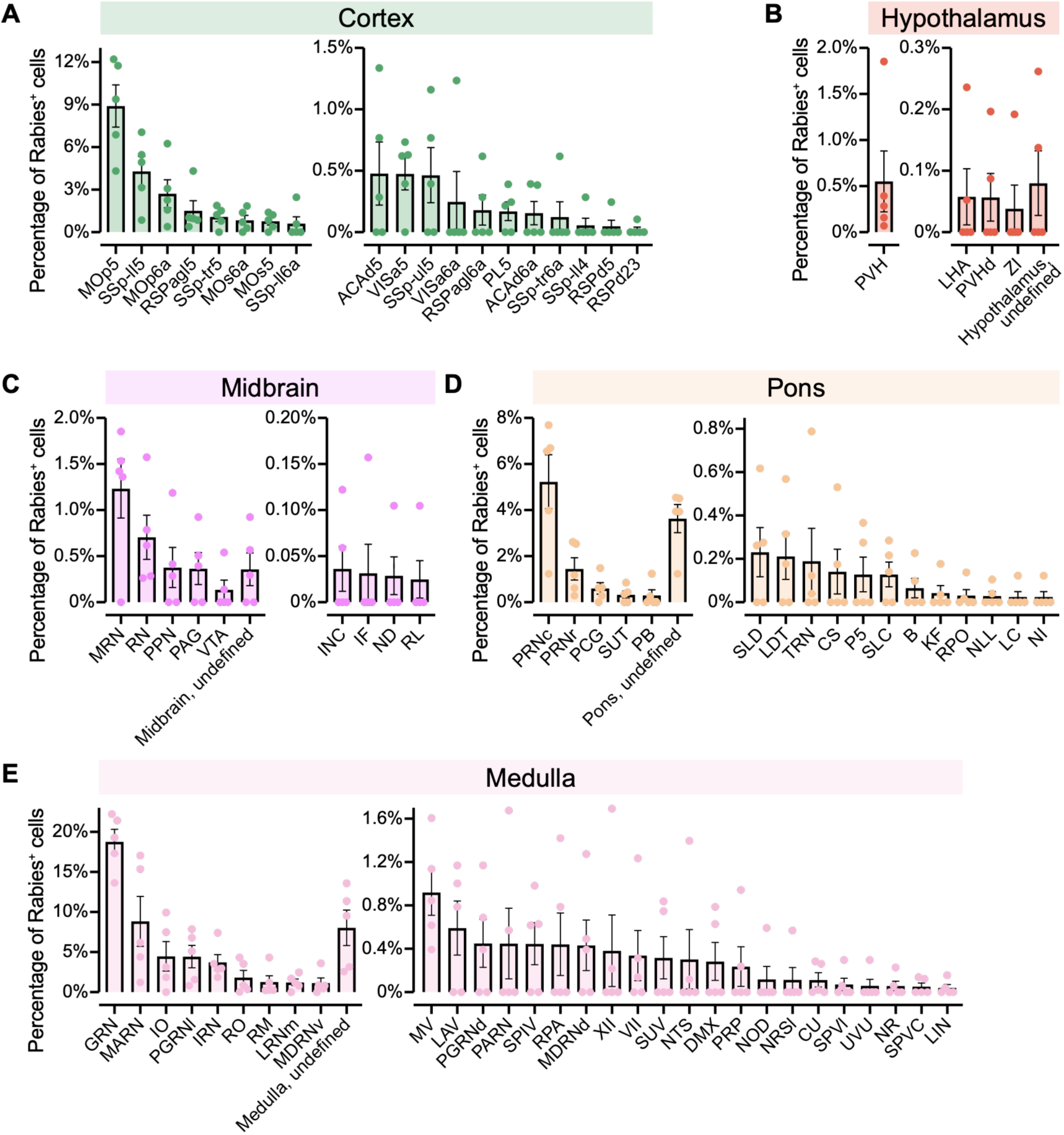
The Celiac-Mesenteric Ganglia (CMG) preganglionic spinal neurons are innervated by neurons residing in brainstem, hypothalamus and cortex. Quantification of rabies^+^ neurons in the 5 brain structures were any rabies^+^ neuron was found. (A) Cortex. MOp5, Primary motor area, layer 5. SSp-ll5, Primary somatosensory area, lower limb, layer 5. MOp6a, Primary motor area, layer 6a. RSPagl5, Retrosplenial area, lateral agranular part, layer 5. SSp-tr5, Primary somatosensory area, trunk, layer 5. MOs6a, Secondary motor area, layer 6a. MOs5, Secondary motor area, layer 5. SSp-ll6a, Primary somatosensory area, lower limb, layer 6a. ACAd5, Anterior cingulate area, dorsal part, layer 5. VISa5, Visual area anterior, layer 5. SSp-ul5, Primary somatosensory area, upper limb, layer 5. VISa6a, Visual area anterior, layer 6a. RSPagl6a, Retrosplenial area, lateral agranular part, layer 6a. PL5, Prelimbic area, layer 5. ACAd6a, Anterior cingulate area, dorsal part, layer 6a. SSp-tr6a, Primary somatosensory area, trunk, layer 6a. SSp-ll4, Primary somatosensory area, lower limb, layer 4. RSPd5, Retrosplenial area, dorsal part, layer 5. RSPd23, Retrosplenial area, dorsal part, layer 2/3. (B) Hypothalamus. PVH, Paraventricular hypothalamic nucleus. LHA, Lateral hypothalamic area. PVHd, Paraventricular hypothalamic nucleus, descending division. ZI, Zona incerta. (C) Midbrain. MRN, Midbrain reticular nucleus. RN, Red nucleus. PPN, Pedunculopontine nucleus. PAG, Periaqueductal gray. VTA, Ventral tegmental area. INC, Interstitial nucleus of Cajal. IF, Interfascicular nucleus raphe. ND, Nucleus of Darkschewitsch. RL, Rostral linear nucleus raphe. (D) Pons. PRNc, Pontine reticular nucleus, caudal part. PRNr, Pontine reticular nucleus, rostral part. PCG, Pontine central gray. SUT, Supratrigeminal nucleus. PB, Parabrachial nucleus. SLD, Sublaterodorsal nucleus. LDT, Laterodorsal tegmental nucleus. TRN, Tegmental reticular nucleus. CS, Superior central nucleus raphe. P5, Peritrigeminal zone. SLC, Subceruleus nucleus. B, Barrington’s nucleus. KF, Kölliker-Fuse subnucleus. RPO, Nucleus Raphe Pontis. NLL, Nucleus of the lateral lemniscus. LC, Locus ceruleus. NI, Nucleus incertus (E) Medulla. GRN, Gigantocellular reticular nucleus. MARN, Magnocellular reticular nucleus. IO, Inferior olivary complex. PGRNl, Paragigantocellular reticular nucleus, lateral part. IRN, Intermediate reticular nucleus. RO, Nucleus raphe obscurus. RM, Nucleus raphe magnus. LRNm, Lateral reticular nucleus, magnocellular part. MDRNv, Medullary reticular nucleus, ventral part. MV, Medial vestibular nucleus. LAV, Lateral vestibular nucleus. PGRNd, Paragigantocellular reticular nucleus, dorsal part. PARN, Parvicellular reticular nucleus. SPIV, Spinal vestibular nucleus. RPA, Nucleus raphe pallidus. MDRNd, Medullary reticular nucleus, dorsal part. XII, Hypoglossal nucleus. VII, Facial motor nucleus. SUV, Superior vestibular nucleus. NTS, Nucleus of the solitary tract. DMX, Dorsal motor nucleus of the vagus nerve. PRP, Nucleus prepositus. NOD, Nodulus. NTSl, Nucleus of the solitary tract, lateral part. CU, Cuneate nucleus. SPVI, Spinal nucleus of the trigeminal, interpolar part. UVU, Uvula. NR, Nucleus of Roller. SPVC, Spinal nucleus of the trigeminal, caudal part. LIN, Linear nucleus of the medulla. N=5 mice.

**Supplemental Video 1**. Whole mount adult cleared spleen and pancreas stained with Tyrosine Hydroxylase (TH), revealing the dense innervation of both organs and its different innervation patterns.

**Supplemental Video 2**. Whole mount Celiac-Mesenteric ganglia from a mouse injected with CTB in the pancreas (green) and spleen (red), revealing the neurons projecting to spleen and pancreas.

**Supplemental Video 3**. Whole mount Nodose ganglia from a mouse injected with CTB in the pancreas (green) and spleen (red), revealing the neurons projecting from spleen and pancreas.

**Supplemental Video 4**. Whole mount Celiac-Mesenteric ganglia from a TH^Cre^;CAG-Sun1/sfGFP mouse shows the localization of TH^+^ nuclei in the mouse line used for SMART-Seq, where GFP^+^ nuclei were sorted for sequencing.

**Supplemental Video 5**. BrainJ reconstruction of the brain indicating all identified rabies^+^ cells within one of the analyzed brains (conf Figure 4).

## REFERENCES

1. Rosas-Ballina, M. et al. Acetylcholine-Synthesizing T Cells Relay Neural Signals in a Vagus Nerve Circuit. Science 334, 98–101 (2011).

2. Zhang, X. et al. Brain control of humoral immune responses amenable to behavioural modulation. Nature 581, 204–208 (2020).

3. Diaz-Salazar, C. et al. Cell-intrinsic adrenergic signaling controls the adaptive NK cell response to viral infection. J. Exp. Med. 217, e20190549 (2020).

4. Globig, A.-M. et al. The β1-adrenergic receptor links sympathetic nerves to T cell exhaustion. Nature 622, 383–392 (2023).

5. Wu, M. et al. Innervation of nociceptor neurons in the spleen promotes germinal center responses and humoral immunity. Cell 187, 2935–2951.e19 (2024).

6. Verlinden, T. J. M. et al. Innervation of the human spleen: A complete hilum-embedding approach. Brain. Behav. Immun. 77, 92–100 (2019).

7. Williams, J. M. et al. Sympathetic innervation of murine thymus and spleen: Evidence for a functional link between the nervous and immune systems. Brain Res. Bull. 6, 83–94 (1981).

8. Romano, T. A., Felten, S. Y., Olschowka, J. A. & Felten, D. L. Noradrenergic and peptidergic innervation of lymphoid organs in the beluga, *Delphinapterus leucas* : An anatomical link between the nervous and immune systems. J. Morphol. 221, 243–259 (1994).

9. Furness, J. B. & Moore, J. The adrenergic innervation of the cardio-vascular system of the lizard Trachysaurus rugosus. Z. F¾r Zellforsch. Mikrosk. Anat. 108, 150–176 (1970).

10. Kinney, K. S., Cohen, N. & Felten, S. Y. Noradrenergic and peptidergic innervation of the amphibian spleen: Comparative studies. Dev. Comp. Immunol. 18, 511–521 (1994).

11. Craig M Flory. Autonomic innervation of the Spleen of the Coho Salmon, Oncorhynchus kisutch: A Histochemical Demonstration and Preliminary Assessment of Its lmmunoregulatory Role.

12. Felten, D. L., Felten, S. Y., Bellinger, D. L. & Madden, K. S. Fundamental aspects of neural-immune signaling. Psychother. Psychosom. 60, 46–56 (1993).

13. Ayers, A. B., Davies, B. N. & Withrington, P. G. Responses of the isolated, perfused human spleen to sympathetic nerve stimulation, catecholamines and polypeptides. Br. J. Pharmacol. 44, 17–30 (1972).

14. Donegà, M., et al. Human-relevant near-organ neuromodulation of the immune system via the splenic nerve. Proc. Natl. Acad. Sci. 118, e2025428118 (2021).

15. Rogausch, H. et al. Endotoxin impedes vasoconstriction in the spleen: role of endogenous interleukin-1 and sympathetic innervation. Am. J. Physiol.-Regul. Integr. Comp. Physiol. 272, R2048–R2054 (1997).

16. Pavlov, V. A. & Tracey, K. J. Neural regulation of immunity: molecular mechanisms and clinical translation. Nat. Neurosci. 20, 156–166 (2017).

17. Bellinger, D. L., Ackerman, K. D., Felten, S. Y. & Felten, D. L. A longitudinal study of age-related loss of noradrenergic nerves and lymphoid cells in the rat spleen. Exp. Neurol. 116, 295–311 (1992).

18. Cleypool, C. G. J. et al. Age-Related Variation in Sympathetic Nerve Distribution in the Human Spleen. Front. Neurosci. 15, 726825 (2021).

19. Lorton, D. et al. Changes in the density and distribution of sympathetic nerves in spleens from Lewis rats with adjuvant-induced arthritis suggest that an injury and sprouting response occurs. J. Comp. Neurol. 489, 260–273 (2005).

20. Hoover, D. B. Cholinergic modulation of the immune system presents new approaches for treating inflammation. Pharmacol. Ther. 179, 1–16 (2017).

21. Koopman, F. A., et al. Vagus nerve stimulation inhibits cytokine production and attenuates disease severity in rheumatoid arthritis. Proc. Natl. Acad. Sci. 113, 8284–8289 (2016).

22. Levine, Y. A., Simon, J., Faltys, M. & Zitnik, R. Bioelectronic Therapy for the Treatment of Rheumatoid Arthritis and Inflammatory Bowel Disease. in Neuromodulation 1503–1511 (Elsevier, 2018). doi:10.1016/B978-0-12-805353-9.00127-3.

23. Natarajan, C., et al. Electrical stimulation of the vagus nerve ameliorates inflammation and disease activity in a rat EAE model of multiple sclerosis. Proc. Natl. Acad. Sci. 121, e2322577121 (2024).

24. Tesser, J. R. P. et al. Vagus nerve-mediated neuroimmune modulation for rheumatoid arthritis: a pivotal randomized controlled trial. Nat. Med. 32, 369–378 (2026).

25. Renier, N. et al. iDISCO: A Simple, Rapid Method to Immunolabel Large Tissue Samples for Volume Imaging. Cell 159, 896–910 (2014).

26. Bratton, B. O. et al. Neural regulation of inflammation: no neural connection from the vagus to splenic sympathetic neurons. Exp. Physiol. 97, 1180–1185 (2012).

27. Sharkey, K. A. & Williams, R. G. Extrinsic innervation of the rat pancreas: Demonstration of vagal sensory neurones in the rat by retrograde tracing. Neurosci. Lett. 42, 131–135 (1983).

28. Principles of Neural Science. (McGraw Hill, New York, 2021).

29. Wang, T., Teng, B., Yao, D. R., Gao, W. & Oka, Y. Organ-specific sympathetic innervation defines visceral functions. Nature 637, 895–902 (2025).

30. Wei, Y. et al. Sympathetic functional units encoded by genetically defined postganglionic neurons. Neuron 114, 463–478.e7 (2026).

31. Ben-Simon, Y. et al. A suite of enhancer AAVs and transgenic mouse lines for genetic access to cortical cell types. Preprint at 10.1101/2024.06.10.597244 (2024).

32. Fiederling, F., Hammond, L. A., Ng, D., Mason, C. & Dodd, J. SpineRacks and SpinalJ for efficient analysis of neurons in a 3D reference atlas of the mouse spinal cord. STAR Protoc. 2, 100897 (2021).

33. Cano, G., Sved, A. F., Rinaman, L., Rabin, B. S. & Card, J. P. Characterization of the central nervous system innervation of the rat spleen using viral transneuronal tracing. J. Comp. Neurol. 439, 1–18 (2001).

34. Harima, Y. et al. Parallel labeled-line organization of sympathetic outflow for selective organ regulation in mice. Nat. Commun. 15, 10478 (2024).

35. Berthoud, H.-R. & Powley, T. L. Characterization of vagal innervation to the rat celiac, suprarenal and mesenteric ganglia. J. Auton. Nerv. Syst. 42, 153–169 (1993).

36. Tracey, K. J. Reflex control of immunity. Nat. Rev. Immunol. 9, 418–428 (2009).

37. Christianson, J. A. et al. Convergence of bladder and colon sensory innervation occurs at the primary afferent level. Pain 128, 235–243 (2007).

38. Torres, H. et al. Sympathetic innervation of the mouse kidney and liver arising from prevertebral ganglia. Am. J. Physiol.-Regul. Integr. Comp. Physiol. 321, R328–R337 (2021).

39. Holmgren, S. Sympathetic innervation of the coeliac artery from a teleost, gadus morhua. Comp. Biochem. Physiol. Part & Comp. Pharmacol. 60, 27–32 (1978).

40. Watson, A. H. D. The structure of the coeliac ganglion of a teleost fish Myoxocephalus scorpius. Cell Tissue Res. 210, (1980).

41. Taxi, J. Morphology of the Autonomic Nervous System. in Frog Neurobiology 93–150 (Springer Berlin Heidelberg, Berlin, Heidelberg, 1976). doi:10.1007/978-3-642-66316-1_3.

42. Du Plessis, M. & Loukas, M. A comprehensive study of the abdominal ganglia part 1: Celiac, phrenic and superior mesenteric ganglia. Clin. Anat. 35, 998–1006 (2022).

43. Vida, G., Peña, G., Deitch, E. A. & Ulloa, L. α7-Cholinergic Receptor Mediates Vagal Induction of Splenic Norepinephrine. J. Immunol. 186, 4340–4346 (2011).

44. Funakoshi, K. & Nakano, M. The Sympathetic Nervous System of Anamniotes. Brain. Behav. Evol. 69, 105–113 (2007).

45. Buijs, R. M., Van Der Vliet, J., Garidou, M.-L., Huitinga, I. & Escobar, C. Spleen Vagal Denervation Inhibits the Production of Antibodies to Circulating Antigens. PLoS ONE 3, e3152 (2008).

46. Vizzard, M. A., Brisson, M. & De Groat, W. C. Transneuronal labeling of neurons in the adult rat central nervous system following inoculation of pseudorabies virus into the colon. Cell Tissue Res. 299, 9–26 (2000).

47. Westerhaus, M. J. & Loewy, A. D. Central representation of the sympathetic nervous system in the cerebral cortex. Brain Res. 903, 117–127 (2001).

48. Dum, R. P., Levinthal, D. J. & Strick, P. L. The mind–body problem: Circuits that link the cerebral cortex to the adrenal medulla. Proc. Natl. Acad. Sci. 116, 26321–26328 (2019).

49. Levinthal, D. J. & Strick, P. L. Multiple areas of the cerebral cortex influence the stomach. Proc. Natl. Acad. Sci. 117, 13078–13083 (2020).

50. Tasic, B. et al. Adult mouse cortical cell taxonomy revealed by single cell transcriptomics. Nat. Neurosci. 19, 335–346 (2016).

51. Tasic, B. et al. Shared and distinct transcriptomic cell types across neocortical areas. Nature 563, 72–78 (2018).

52. Korsunsky, I. et al. Fast, sensitive and accurate integration of single-cell data with Harmony. Nat. Methods 16, 1289–1296 (2019).

53. Gayoso, A. et al. A Python library for probabilistic analysis of single-cell omics data. Nat. Biotechnol. 40, 163–166 (2022).

54. Luecken, M. D. et al. Benchmarking atlas-level data integration in single-cell genomics. Nat. Methods 19, 41–50 (2022).

